# Neuronal DNA repair mediates systemic mitochondrial maintenance through acetylcholine signaling

**DOI:** 10.64898/2026.02.11.705232

**Authors:** Paulo F. L. da Silva, Baptiste Ropert, Matthias Rieckher, Corvin Rive, Björn Schumacher

## Abstract

DNA repair defects such as Nucleotide Excision Repair (NER) mutations can lead to progeroid syndromes characterized by growth retardation and accelerated aging with distinct tissue type specific pathologies. DNA damage induced dysfunction of specific cell types can impair the organism’s overall function. It is unknown whether, in reverse, DNA repair in specific tissues can have non-cell-autonomous consequences on other tissues carrying DNA damage. To explore the organismal consequences of cell type specific DNA repair, we restricted NER activity to specific tissues and monitored consequences on organismal health in *C. elegans.* Unexpectedly, we observed that tissue specific DNA repair is sufficient to improve overall DNA damage resistance. Strikingly, the sole NER activity in neurons is sufficient to provide similar overall DNA damage resistance as whole body NER. We show that this systemic DNA damage resilience is mediated by neuronal Acetylcholine signaling that promotes the maintenance of mitochondrial activity throughout the organism.

## Introduction

The genome is constantly exposed to genotoxic threats stemming from the normal cellular metabolism^1^ and due to external sources^2^ leading to genome instability that causes cancer^3^ and promotes aging^4,5^. Among the specialized pathways to remove the DNA damage ^6^, Nucleotide Excision Repair (NER) is a versatile mechanism that removes bulky lesion types that lead to the stalling of RNA and DNA polymerases. UV-induced cyclobutane pyrimidine dimers (CPDs) and 6-4 photoproducts are paradigm damage types that lead to a distortion of the DNA double-helix. In humans, defects in global genome (GG)-NER, caused by mutations in XPC, XPA, the TFIIH-XPB-XPD complex or in the XPF and XPG endonucleases, lead to Xeroderma Pigmentosum (XP). XP patients display photosensitivity, 1000-fold, increased risk of developing cancer in the skin and eye, and in approximately 30% patients, including those with XPA mutations, progressive neurological degeneration^7–9^. Defects in transcription-coupled (TC)-NER activity, caused by mutations in either the lesion recognition factors CSA or CSB lead to Cockayne’s syndrome (CS). CS patients display accelerated ageing phenotypes in multiple tissues, including severe neurodegeneration, hearing loss, cataracts and cachexia, as well as growth impairment^10^. Additionally, point mutations in XPB, XPD, and TTDA are also associated with Trichothiodystrophy (TTD), a progeroid syndrome that very much resembles CS but in which patients also display brittle hair and nails^11^.

The heterogenous outcomes of NER deficiency highlight the segmental consequences of DNA damage accumulation, i.e. different tissues are differentially affected. This suggests that the requirement for the DNA repair machinery is not necessarily similar in all cell types. However, it is still unclear whether distinct tissues employ different strategies to maintain functionality and whether more complex inter-tissue signaling responses are involved in the maintenance of tissue functionality when faced with DNA damage.

XPA is a key NER factor involved in damage verification, DNA strand unwinding and assembly of the endonuclease incision complex^12–14^. Due to its capacity to bind proteins involved in every step of the NER pathway, as well as other regulatory factors, such as ATR and PARP-1^15^, XPA plays a central role in NER regulation. In *C. elegans*, *xpa-1* loss-of-function mutations lead to hypersensitivity to UV-induced DNA damage during development and adulthood ^16–19^.

Here, we generated a model that allows testing the requirement of cell type specific DNA repair using *C. elegans* as organismal model featuring specialized neuronal, intestinal and muscle cells. We employed UV irradiation as it leads to specific and well-defined lesions that strictly require NER for their removal. To generate an experimental model with cell type restricted DNA repair activity, we employed *xpa-1(ok698)* mutant animals that are completely NER deficient and reintroduced NER capacity by transgenic expression of XPA-1::GFP under the control of endogenous, neuron, muscle, or intestine specific promoters. We show that the tissue-specific restriction of NER is sufficient to partially protect the animals from DNA damage. Most strikingly, restricting NER only to neurons conferred a similar level of protection against DNA damage as observed in animals with the endogenous reconstitution of NER. Neuronal DNA repair was sufficient to reverse DNA damage-induced pathologies including developmental growth retardation and accelerated aging and maintain health parameters with the exception of intestinal function. We further determined that the systemic effect of *xpa-1* relies on the Acetylcholine signalling pathway, unravelling a previously unknown function of this neurotransmitter in regulating the DNA damage response. We further observed that the neuron-specific NER activity reversed the systemic dampening of mitochondrial activity, which we propose is sufficient for mitigating the organismal consequences of DNA damage.

## Results

### DNA repair restoration by tissue-specific XPA-1::GFP expression

In order to determine the tissues where XPA-1 is expressed in normal conditions, we generated a transgenic *C. elegans* strain expressing XPA-1 C-terminally-tagged with GFP under the endogenous *xpa-1* promoter. We observed XPA-1 expression throughout all developmental stages, with very high expression in the embryo, and during adulthood in several somatic tissues, including the intestine, muscle, cuticle, and neurons (**Figure 1A**). To better understand the contribution of tissue-specific *xpa-1* activity in organismal responses to DNA damage, we generated transgenic lines that express XPA-1 fused to GFP in specific tissues (**Supplementary Table 1**). We stably integrated the XPA-1::GFP transgene under control of distinct tissue-specific promoters in a *xpa-1* null background (*ok(698)* allele), in which approximately half of the ORF is deleted^20,21^, thus generating tissue-specific *xpa-1* rescue lines. Following this strategy, we developed four rescue lines: an endogenous *xpa-1* rescue line (Endogenous rescue) under control of the endogenous *xpa-1* promoter (*xpa-1(ok698)*;[*p_xpa-1_::gfp::xpa-1;p_myo-2_::tdTomato*]), an intestinal Rescue line (Intestinal rescue) under control of the *ges-1* promoter (*xpa-1(ok698)*;[*p_ges-1_::gfp::xpa-1;p_myo-2_::tdTomato*]), a muscle Rescue line (Muscle rescue) under control of the *myo-3* promoter (*xpa-1(ok698)*;[*p_myo-3_::gfp::xpa-1;p_myo-2_::tdTomato*]), and a neuronal Rescue line (Neuronal rescue), under control of the pan-neuronal *rgef-1* promoter (*xpa-1(ok698)*;[*p_rgef-1_::gfp::xpa-1;p_unc-122_::gfp*]). Using confocal microscopy, we observed GFP fluorescence exclusively in the targeted tissues in each rescue line, thus confirming the tissue-specific expression of XPA-1::GFP (**Figures 1B – E**).

**Figure 1.**
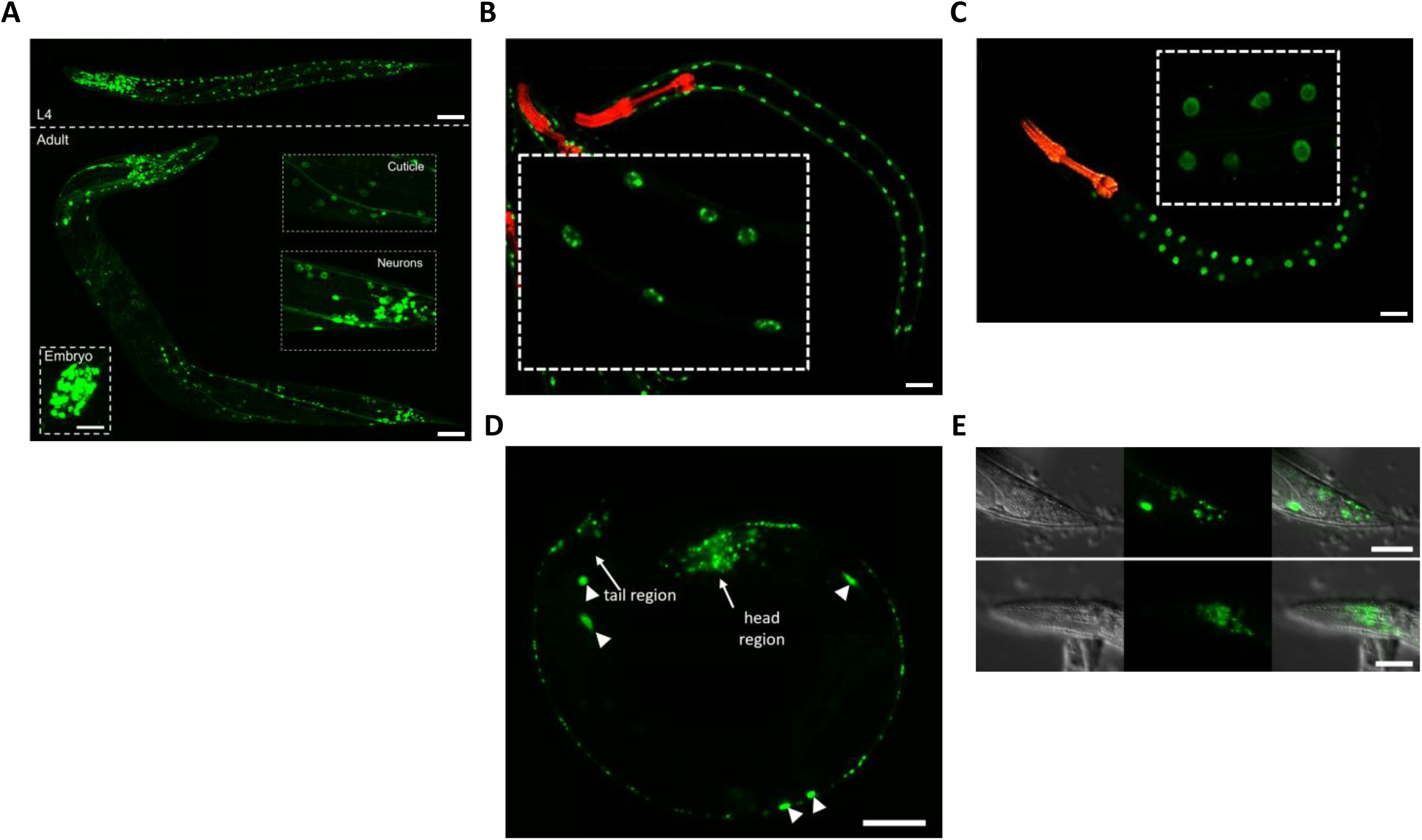
Tissue-specific XPA-1::GFP expression in *C. elegans*. (**A**) Representative fluorescence images of L4 (top panel), adult (bottom panel) and embryo (bottom left) from nematodes showing XPA-1::GFP expression under the endogenous *xpa-1* promoter. Cuticle and neuronal system displaying particularly clear expression are highlighted in right boxes. Scale bars = 50µm. (**B**) Representative fluorescence images of XPA-1::GFP expression in the nuclei of intestinal cells (*pges-1::xpa-1::gfp*) and tdTomato expression in the pharynx (*pmyo-2::tdTomato*) (co-injection marker), from day 1 adults. Scale bar = 50µm. (**C**) Representative fluorescence images of XPA-1::GFP expression in the nuclei of muscle cells (*pmyo-3::xpa-1::gfp*) and tdTomato expression in the pharynx (*pmyo-2::tdTomato*) (co-injection marker)), from day 1 adults. Scale bar = 50µm. (**D**) Representative fluorescence images of XPA-1::GFP expression in the nuclei of neuronal cells (*prgef-1::xpa-1::gfp*), And GFP expression in coelomocytes (*punc-122::gfp*) (co-injection marker) (white triangles), from day 1 adults. Scale bar = 50µm. (**E**) Representative fluorescence images of XPA-1::GFP expression in the nuclei of tail (top panel) and head (bottom panel) neurons (*prgef-1::xpa-1::gfp*), from day 1 adults. Scale bar = 50µm.

### Neuron-specific NER activity is sufficient for organismal DNA damage resistance

To evaluate how tissue-specific DNA repair influences the organism’s DNA damage sensitivity, we analyzed the delay in developmental growth in the tissue-specific XPA-1 rescue lines after UVB exposure at L1 larvae stage^22,23^. 72 h post UV treatment of the L1 larvae, we observed some rescue effect of the UV hypersensitivity of *xpa-1* mutants in either of the rescue strains **(Figure 2A)**. We then quantified the larval development through the L2, L3, and L4 stages until adulthood. As expected^17,18^, in contrast to wild-type animals, *xpa-1(ok698)* failed to fully develop to adults upon 4 mJ/cm² UV irradiation. The *xpa-1* mutants carrying the transgenic XPA-1::GFP under the endogenous promoters (Endogenous rescue) showed a full restoration of developmental progression at low UV-doses and mild delay at higher doses, with a small decrease in fully developed worms upon 8 mJ/cm² UV. The observed significant yet partial rescue is typically observed in transgenic rescue lines and confirms the efficiency of the rescue transgene to compensate for the mutated *xpa-1* gene. Interestingly, each of the tissue-specific rescue lines alleviated the UV sensitivity in the *xpa-1* mutant background, however, to different degrees. The Muscle rescue showed the least efficient reversal of the UV sensitivity, with a significant decrease of fully developed worms upon 4 mJ/cm² UVB. The Intestinal rescue, showed a more pronounced rescue of the developmental growth defect with a significant but milder decrease of fully developed worms upon 4 mJ/cm² UVB. The Neuronal rescue showed the most efficient rescue of the UV sensitivity that was almost comparable with the Endogenous rescue, with a significant decrease of fully developed worms only upon 8 mJ/cm² **(Figure 2B)**.

**Figure 2.**
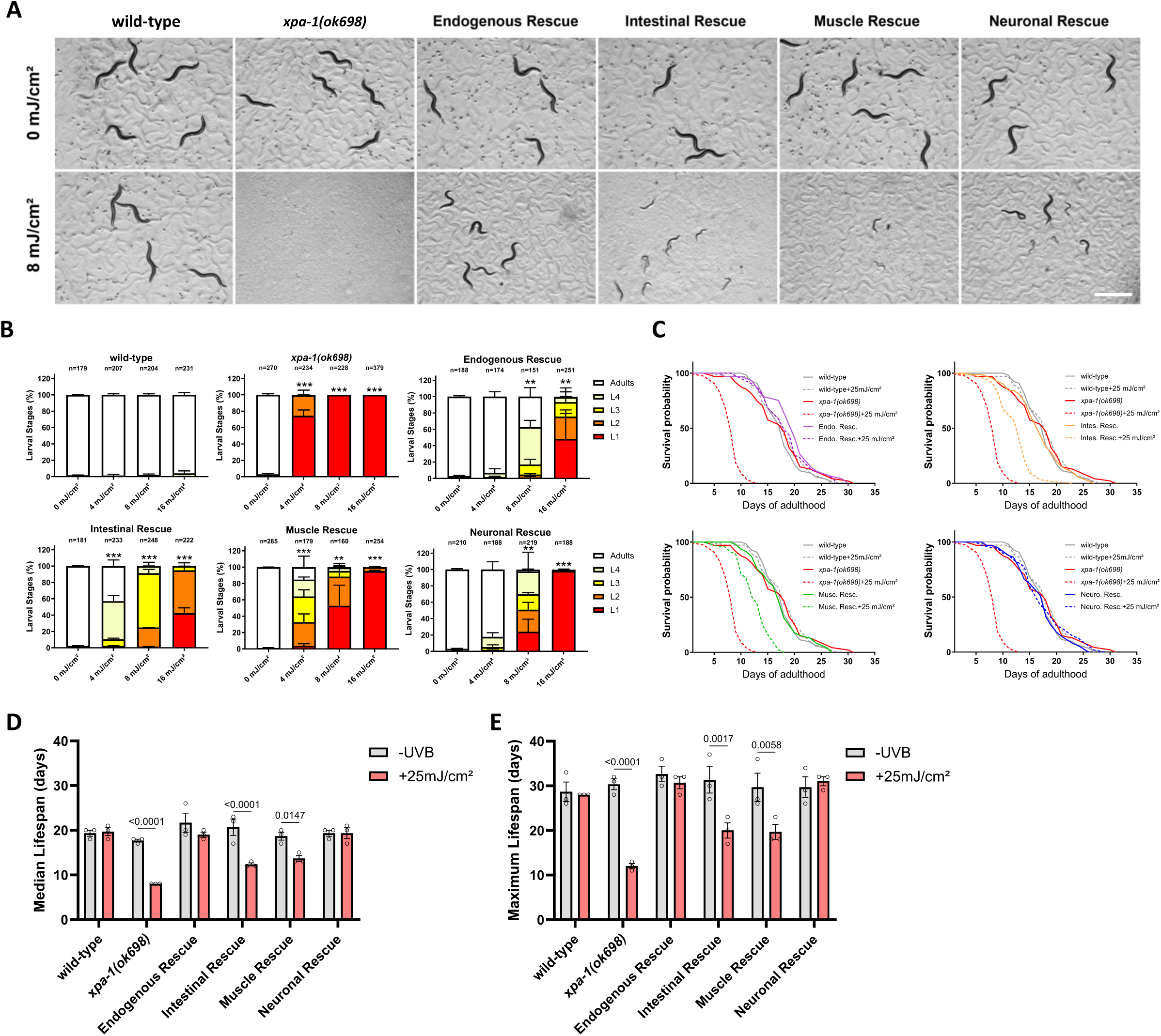
Tissue-specific XPA-1 expression differentially rescues *xpa-1* UV-induced developmental arrest and lifespan shortening. (**A**) Representative pictures of irradiated (8mJ/cm²) and non-irradiated wild-type, *xpa-1* and *xpa-1* tissue-specific rescue lines, 72 hours after UVB irradiation at L1 stage. Scale Bar = 1mm. (**B**) Quantification of development of wild-type, *xpa-1* and *xpa-1* tissue-specific rescue lines, 72 hours after UVB irradiation at L1 stage. Data represent Mean+SD of N=3 biological replicates. Statistical analysis: ANOVA. *p*-value:*<0.05, **<0.01, ***<0.001. (**C**) Lifespan assays of wild-type, *xpa-1* and *xpa-1* tissue-specific rescue lines after UVB irradiation at day 1 adult stage. Representative survival curve from one individual experiment. n>100 per condition. (**D**) Quantification of median lifespan and (**E**) maximum lifespan from survival curves displayed in C. Data represent Mean+SD of N=3 biological replicates. Statistical analysis: Two-way ANOVA followed by Post-hoc Tukey.

In order to independently verify the effect of cell type-specific NER restriction, we performed the same analysis on strains carrying the transgenes in the form of extrachromosomal arrays. When we treated those strains with UVB, we observed a most striking rescue when driving XPA-1 expression either by the endogenous *xpa-1* promoter as well as the neuron-and intestine-specific promoters and a less efficient rescue in muscle-specific XPA-1 expression in the *xpa-1* mutant background (**Figure S1A**).

Exposure to genotoxic insults in adult animals is known to reduce lifespan. NER-deficient mutant worms in particular, show a robust lifespan reduction when exposed to UVB light^23^. To determine whether rescuing DNA repair in specific tissues can rescue the lifespan reduction of *xpa-1* mutants, we carried out lifespan assays using both the integrated tissue-specific **(Figure 2C)** and the extrachromosomal **(Figure S1B)** *xpa-1* rescue lines. In accordance with previous results, *xpa-1(ok698)* showed a drastic reduction of both median and maximum lifespan (-55% and-60% respectively) at a mild UV dose that was well tolerated by wild-type animals. The lifespan of Endogenous rescue animals was not decreased upon genotoxic stress contrary to the Intestinal rescue and Muscle rescue strains, whose median and maximum lifespan were significantly decreased but with a milder effect compared to *xpa-1(ok698)* (Intestinal rescue:-40% and-36%; Muscle rescue:-27% and-34%). Strikingly, the Neuronal rescue showed no decrease in lifespan compared to control and Endogenous rescue worms **(Figure 1D-E)**.

Altogether, these data indicate that NER activity in either intestine, muscle, or neurons can contribute to the organism’s resistance to UV-induced DNA damage. Moreover, neuron-specific NER activity is nearly as effective in conferring DNA damage resistance during development and aging as endogenously restored NER activity.

### Neuron-specific DNA repair exerts systemic effects

To establish how cell type specific restrictions of NER activity affect physiological parameters of health, we measured the integrity of muscles, neurons and the gastrointestinal tract after UV exposure in each of the tissue-specific rescue strains. To observe the muscles structure, we stained the animals with Phalloidin 24 hours after UV irradiation. We observed that, in contrast to wild-type, the muscle structure in *xpa-1(ok698)* mutants was altered after UV exposure similarly to the Intestinal rescue strain. Both Endogenous rescue, Muscle rescue and, surprisingly, the Neuronal rescue strain retained the muscle structure after UV in similar fashion to wild-type animals **(Figure 3A).** To quantify muscle capacity, we analyzed the average and maximum speed of each strain with or without DNA damage induction. Motility features of *xpa-1(ok698)*, Intestinal rescue, Muscle rescue and Neuronal rescue show that these worms are moving slower than the wild-type and Endogenous rescue strains. Upon DNA damage induction, we observed a significant decrease in motility in *xpa-1(ok698)* (Average speed:-81%; Max speed:-64%) and in Intestinal rescue (Average speed:-74%; Max speed:-58%) but not in Endogenous rescue, Muscle rescue and Neuronal rescue **(Figure 3B-C)**. Pharyngeal pumping necessitates a functioning neuromuscular axis and is indicative of muscular function and overall health but also survival probability^24,25^. Except for wild-type and Endogenous rescue strains, all strains displayed a significant reduction in pumping frequency upon DNA damage. While the *xpa-1(ok698)* mutants and the Intestinal rescue had a very strong reduction (-88% and-67% respectively), Muscle rescue and Neuronal rescue displayed only a mild decreased in pumping frequency (-18% and-17% respectively) **(Figure 3D)**. These results suggest that DNA repair is required for maintaining somatic functions particularly in the neuromuscular axis, as a rescue of *xpa-1* expression in either muscle or neuron is sufficient to restore muscle function amid DNA damage.

**Figure 3.**
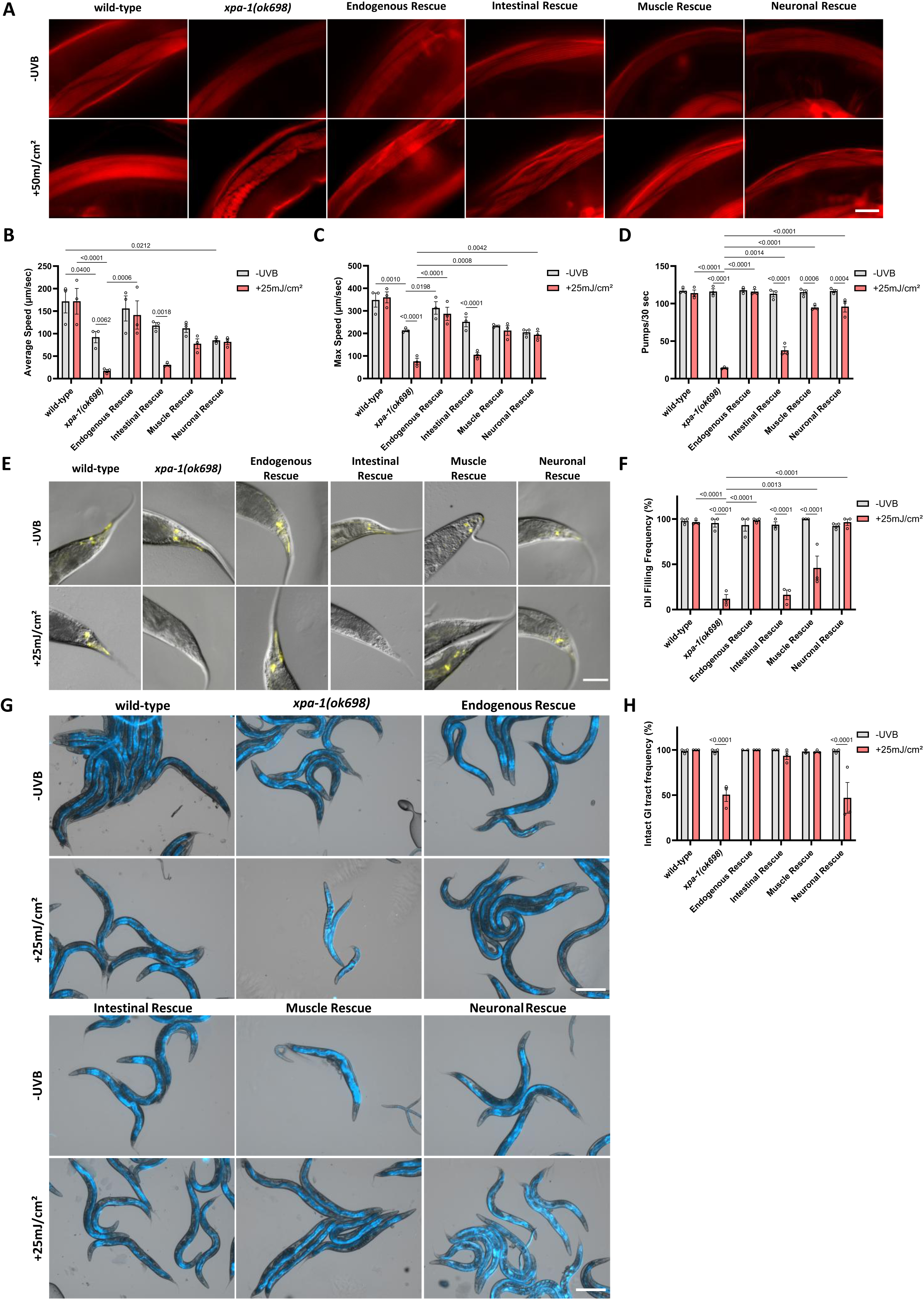
Neuron-specific XPA-1 expression rescues UV-induced phenotypes in other tissues. (**A**) Phalloidin staining of muscle cells from day 1 adults from wild-type, *xpa-1(ok698)* and *xpa-1* tissue-specific rescue lines, 24 hours after UVB irradiation. Scale bar = 10µm. (**B**) Average and (**C**) maximum speed measurements from day 1 adults from wild-type, *xpa-1(ok698)* and *xpa-1* tissue-specific rescue lines, 24 hours after UVB irradiation. Data represent Mean+SD of N=3 biological replicates. Statistical analysis: Two-way ANOVA followed by Post-hoc Tukey. (**D**) Quantification of pharyngeal pumping rate from day 1 adults from wild-type, *xpa-1(ok698)* and *xpa-1* tissue-specific rescue lines, 24 hours after UVB irradiation. Data represent Mean+SD of N=3 biological replicates. Statistical analysis: Two-way ANOVA followed by Post-hoc Tukey. (**E**) Representative pictures of DiI staining of Phasmid neurons from day 1 adults from wild-type, *xpa-1(ok698)* and *xpa-1* tissue-specific rescue lines, 24 hours after UVB irradiation. Scale bar = 15µm. **(F)** Quantification of worms harboring intact phasmid neurons in day 1 adults wild-type, *xpa-1(ok698)* and *xpa-1* tissue-specific rescue lines, 24 hours after UVB irradiation. Data represent Mean+SD of N=3 biological replicates. Statistical analysis: Two-way ANOVA followed by Post-hoc Tukey. (**E**) Representative picture of Brilliant Blue staining of intestinal tracts from day 1 adults from wild-type, *xpa-1(ok698)* and *xpa-1* tissue-specific rescue lines, 24 hours after UVB irradiation. Scale bar = 50µm. **(F)** Quantification of worms harboring intact intestinal tract in day 1 adults wild-type, *xpa-1(ok698)* and *xpa-1* tissue-specific rescue lines, 24 hours after UVB irradiation. Data represent Mean+SD of N=3 biological replicates. Statistical analysis: Two-way ANOVA followed by Post-hoc Tukey.

To assess neuronal integrity, we took advantage of the Amphid and Phasmid sensory neurons that can be stained using 1,1’-Dioctadecyl-3,3,3’,3’-Tetramethylindocarbocyanine Perchlorate (DiI) dye when intact. Upon breakage of the axon, the neurons cannot uptake the dye and are not visible^26,27^. We observed a massive decrease in the DiI-positive neurons after UV exposure in *xpa-1(ok698)* mutant (-95%) compared to wild-type animals, indicative of a strong neuronal DNA damage sensitivity. In contrast, no decrease in DiI positive neurons was observed in Endogenous rescue and Neuronal rescue. The decrease in DiI positive neurons frequency in Intestinal rescue was comparable to *xpa-1(ok698)* (-82%) and intermediary in Muscle rescue (-54%) **(Figure 3E-F)**.

We next evaluated the gastrointestinal (GI) tract using Brilliant Blue FCF staining (“Smurf assay”). When mixed with food, this dye is ingested by the animals and stains the intact GI tract. Upon GI tract loss of structural integrity, the dye passes through the intestinal barrier into other tissues and stains the whole animal blue^28^. Contrary to all other tests we performed in the different tissue-specific *xpa-1* rescue strains, only the Neuronal rescue strain was not able to maintain GI tract intact after UVB exposure and showed significantly decreased levels of intact GI tracts, reaching levels comparable to *xpa-1(ok698)* (-52% and-49% respectively); All other strains showed no signs of sensitivity to DNA damage in this assay **(Figure 3G-H)**.

Here, our results show that DNA repair in the neurons alone is sufficient to maintain the animals’ overall health with the exception of the GI tract. Amid neuron-restricted repair, particularly the neuromuscular function remained intact suggesting a non-cell-autonomous effect of neuronal NER activity.

### Acetylcholine signaling is required for systemic DNA damage resistance

To better understand how DNA repair in neurons coordinates maintenance of tissue functionality peripherally, we tested whether specific types of neuronal signaling, potentially involved in inter-tissue communication, influence organismal sensitivity to DNA damage. For this, we performed developmental assays using two mutant strains: *unc-13(s69)* and *unc-31(e928)*, deficient in key synaptic vesicle priming/fusion proteins essential for small-clear vesicle (SCV) and dense-core vesicle (DCV) exocytosis, respectively^29–31^. SCVs are employed to secrete neurotransmitters such as glutamate, γ-Aminobutyric acid (GABA) and acetylcholine (ACh), while DCVs contain larger neuropeptides and monoamines^31^ **(Figure 4A)**. Both *unc-13(s69)* and *unc-31(e928)* mutants possess normal nervous system architecture but an uncoordinated (unc) phenotype^29,31^. Upon UV-induced DNA damage, the *unc-13(s69)* mutants displayed a significant developmental growth delay compared to wild-type animals, with a significant reduction in fully developed animals upon 25 mJ/cm² UVB. On the contrary, *unc-31(e928)* mutants showed no sensitivity to DNA damage, similar to wild-type animals, pointing toward a role of neurotransmitter but not neuropeptide signaling in the response to DNA damage **(Figure 4B)**.

**Figure 4.**
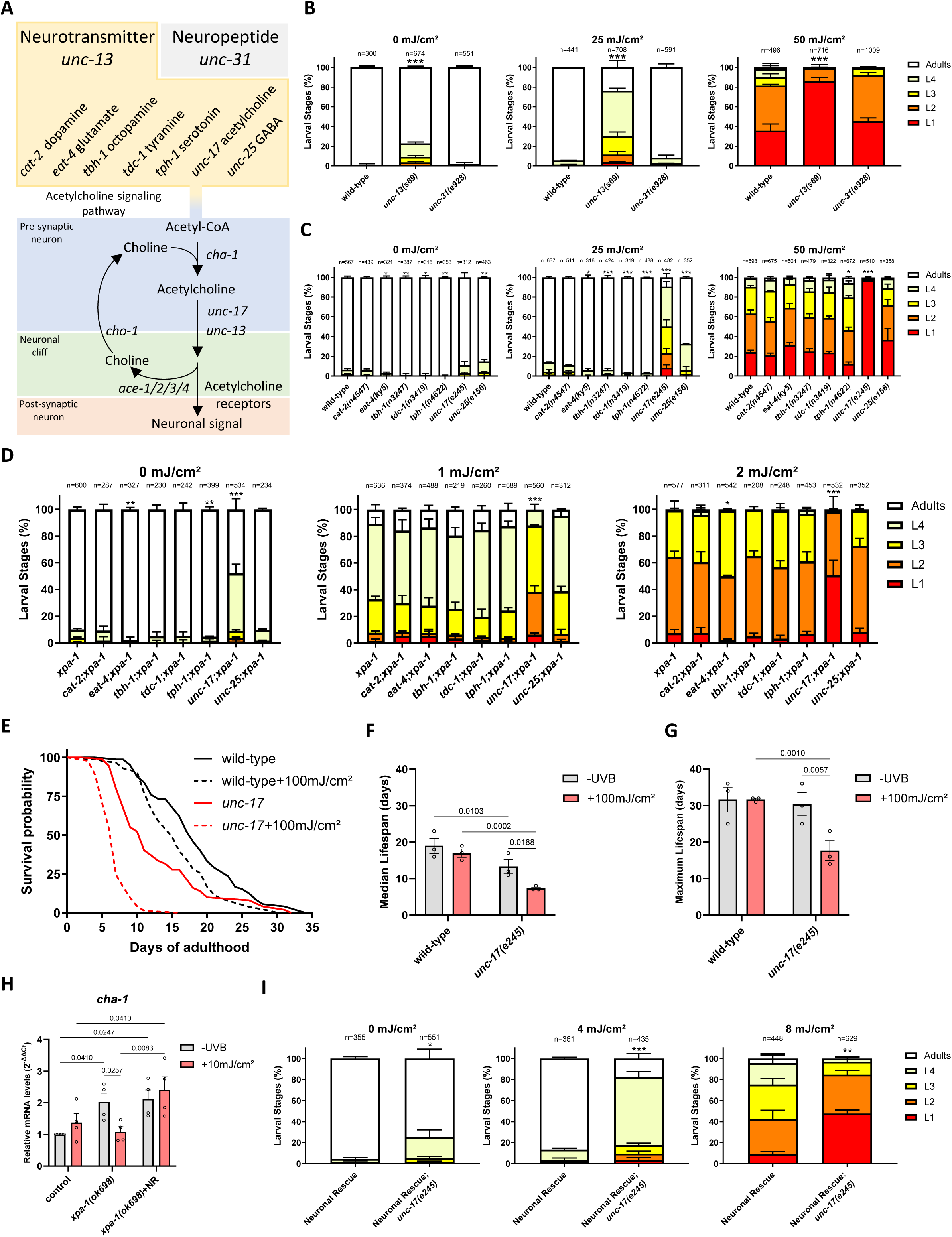
Loss of acetylcholine signaling abolishes XPA-1 Neuronal-rescue effects. **(A)** Scheme of Acetylcholine signaling pathway. **(B)** Quantification of development of wild-type, neuropeptide deficient (*unc-13(s69)*) and neurotransmitter deficient (unc-31*(e928)*) lines, 72 hours after UVB irradiation at L1 stage. Data represent Mean+SD of N=3 biological replicates. Statistical analysis: ANOVA. *p*-value:*<0.05, **<0.01, ***<0.001. **(C)** Quantification of development of wild-type, and major neurotransmitter pathways mutants lines (dopamine (*cat-2(n4547)*, glutamate (*eat-4(ky5)*), octopamine (*tbh-1(n3247)*), tyramine (*tdc-1(n3419)*), serotonin (*tph-1(n4622)*), acetylcholine (*unc-17(e245)*), GABA (*unc-25(e156)*), 72 hours after UVB irradiation at L1 stage. Data represent Mean+SD of N=3 biological replicates. Statistical analysis: ANOVA. *p*-value:*<0.05, **<0.01, ***<0.001. **(D)** Quantification of development of *xpa-1(ok698) and xpa-1(ok698)* mutant worms crossed with major neurotransmitter pathways mutant worms from (C), 72 hours after UVB irradiation at L1 stage (for clarity, only the name of the mutated genes are displayed). Data represent Mean+SD of N=3 biological replicates. Statistical analysis: ANOVA. *p*-value:*<0.05, **<0.01, ***<0.001. (**E**) Lifespan assays of wild-type and acetylcholine signaling mutant (*unc-17(e245)*) lines after UVB irradiation at day 1 adult stage. Representative survival curve from one individual experiment. n>100 per condition. (**F**) Quantification of median lifespan and (**G**) maximum lifespan from survival curves displayed in (E). Data represent Mean+SD of N=3 biological replicates. Statistical analysis: Two-way ANOVA followed by Post-hoc Tukey. **(H)** Quantification of *cha-1* mRNA levels in wild-type, *xpa-1(ok698)* and Neuronal rescue strains, 24 hours after UVB irradiation. Data represents Mean+SD of N=4 biological replicates. Statistical analysis: Two-way ANOVA followed by Post-hoc Tukey. **(I)** Quantification of development of *xpa-1(ok698) and xpa-1(ok698); unc-17(e245)* mutant strains, 72 hours after UVB irradiation at L1 stage. Data represent Mean+SD of N=3 biological replicates. Statistical analysis: ANOVA. *p*-value:*<0.05, **<0.01, ***<0.001.

To determine whether specific neurotransmitters are required for responding to DNA damage, we performed developmental assays after UV exposure using multiple loss-of-function/null mutant strains with defects in all major neurotransmitter signaling pathways: dopamine (*cat-2(n4547)*), glutamate (*eat-4(ky5)*), octopamine (*tbh-1(n3247)*), tyramine (*tdc-1(n3419)*), serotonin (*tph-1(n4622)*), ACh (*unc-17(e245)*) and GABA (*unc-25(e156)*) signaling pathways **(Figure 4A)**. Out of the seven mutants, only the *unc-17(e245)* mutant, coding for the *C. elegans* ortholog of the human vesicular Acetycholine transporter (*VAChT*)^32^, showed an increased DNA damage sensitivity, with a significant decrease of fully developed adult worms upon 25 mJ/cm² UV similarly to the *unc-13(s69)* mutants **(Figure 4C)**, indicating a specific role of ACh signaling in the response to DNA damage. We next tested genetic epistasis between NER and ACh signaling. When exposed to UV, *unc-17(e245); xpa-1(ok698)* double mutant animals showed an increased sensitivity compared to either *unc-17(e245)* or *xpa-1(ok698)* single mutants indicating that ACh signaling is required for DNA damage resistance independently of the NER activity itself **(Figure 4D).**

To determine whether abrogation of ACh signaling also facilitates the onset of somatic decline upon DNA damage induction, we assessed the lifespan of *unc-17(e245)* mutants **(Figure 4E)**. *unc-17(e245)* worms display a robust reduction in median (-30%), but not maximum, lifespan even under basal conditions. Both median and maximum lifespan (-45% and-42% respectively) were severely reduced following exposure to 100 mJ/cm² UV radiation, a dose that does not significantly impact wild-type lifespan **(Figure 4F-G)**. These results further support that *unc-17(e245)* mutant animals display DNA damage sensitivity, indicating that ACh signaling regulates *C. elegans* lifespan, both under normal conditions and following DNA damage.

To confirm that impaired ACh signaling is indeed responsible for increased DNA damage sensitivity, we evaluated the UV sensitivity of mutants defective in other components of the cholinergic system i.e. in addition to *unc-17* and *unc-13*, *cha-1*, *cho-1* and *ace-1/2/3/4*. *cha-1* encodes for the choline acetyltransferase that synthetizes ACh using choline and Acetyl-CoA^33,34^. Following this step, ACh is then packaged into vesicles by VAChT ^35^ and released in SCVs into the synaptic cleft. In the synaptic cleft, ACh is degraded into choline by Acetylcholine esterases encoded by *ace-1/2/3/4*^33,34^. Finally, choline can be reuptaken into the presynaptic neuron via the choline transporters encoded by *cho-1*^33,34^ **(Figure 4A)**. When exposed to UV irradiation as L1 larvae, both *cha-1(p503)* **(Figure S2A)** and *cho-1(tm373)* **(Figure S2B)** showed increased DNA damage sensitivity as the developmental delay was more pronounced than in wild-type animals. Their sensitivity was lower than *unc-17(e245)* and *unc-13(s69)* as the decrease of fully developed animals appeared upon 50 mJ/cm² UVB. In contrast, *ace-2(g72);ace-1(p1000)* mutants, where ACh is stabilized, display resistance to DNA damage as the developmental arrest was mildly decreased **(Figure S2C)**. These results further establish the involvement of ACh signaling in the response to DNA damage. However, ACh supplementation failed to alleviate the developmental delay in *unc-17(e245)* mutants **(Figure S2D)**, *xpa-1(ok698)* or the Neuronal rescue strain **(Figure S2E)**. It is, however, conceivable that the extrinsic ACh supplementation does not restore the lack of ACh release at neuronal synapsis.

In order to assess whether the production of ACh might be impacted in the *xpa-1(ok698)* or the Neuronal rescue strains, we assessed the mRNA levels of the rate-limiting enzyme of the ACh signaling system, encoded by the *cha-1* gene. We observed that *cha-1* expression levels are significantly up-regulated in both *xpa-1(ok698)* (+102%) and the Neuronal rescue strains (+111%) when compared to wild-type (**Figure 4H**). Given that ACh deficiency alone leads to developmental growth delays in NER deficient animals as evidenced by the developmental growth delay of *unc-17(e245);xpa-1(ok698)* double mutants even without UV irradiation (**Figure 4D**), the strong induction of *cha-1* in those two strains suggests that increased levels of ACh are required amid DNA repair deficiency. Upon UVB irradiation, we observed a significant decrease of *cha-1* mRNA levels (-54%) in the *xpa-1(ok698)* strain while they remained unchanged in the wild-type and Neuronal rescue strains (Figure 4H). This suggests that the neuronal DNA repair allows the maintenance of high ACh levels after DNA damage induction.

Next, we aimed to verify whether the DNA damage resistance observed in the Neuronal rescue strain was indeed mediated by ACh signaling. We generated the double mutant transgenic strain *unc-17(e245);xpa-1(ok698)*;[*p_rgef-1_::gfp::xpa-1;p_unc-122_::gfp*] (i.e. Neuronal rescue; *unc-17(e245)*) and assessed sensitivity to DNA damage during development of these animals whose NER activity is restricted to the neurons and whose ACh signaling is blunted. We observed that with UVB doses that allow almost all Neuronal rescue animals to reach adulthood (4 mJ/cm²), the Neuronal rescue; *unc-17(e245)* strain showed a significantly reduced proportion of fully developed adult worms, and at higher dose (8 mJ/cm²) the UV resistance of Neuronal rescue animals was significantly abolished by the *unc-17(e245)* mutation **(Figure 4I)**. These results indicate that the organism’s DNA damage resistance mediated by neuron-specific DNA repair requires ACh signaling.

### Neuronal DNA repair maintains systemic mitochondrial function via ACh signaling

Mitochondrial dysfunction was observed in *xpa-1* mutant *C. elegans* and mice and suggested to result from the deprivation of NAD+ pools amid nuclear DNA damage (as NER has been established to strictly operate in the nucleus but is absent in mitochondria)^36^. ACh has been implicated in stress resistance signaling pathways in cardiomyocytes by regulating mitochondrial dynamics/function^37–39^. We thus hypothesized that ACh signaling could impact the animal’s DNA damage sensitivity by boosting mitochondrial activity. Accordingly, disruption of ACh signaling would result in systemic mitochondrial dysfunction upon DNA damage. To test this hypothesis, we analyzed mitochondrial activity in the different neuronal strains as well as in *xpa-1(ok698)* mutants. First, we used Tetramethylrhodamin-Ethylester (TMRE) staining to measure the mitochondrial membrane potential (MMP), a commonly used proxy for mitochondrial health^40,41^. We analyzed the effects of DNA damage induction in neuropeptide (*unc-13(s69)*) and neurotransmitter (*unc-31(e928)*) deficient strains. In line with their respective sensitivity to DNA damage **(Figure 3B)**, animals lacking a functional neurotransmitter pathway showed strongly reduced MMP levels (-85%), while the lack of a functional neuropeptide pathway had no effect **(Figure 5A)**. Moreover, ACh deficient *unc-17(e245)* mutants showed decreased MMP levels in non-DNA damage condition (-49 %) that is further exacerbated upon DNA damage (-75 %) **(Figure 5B)**. These results show that mitochondrial activity is impaired in ACh signaling mutants, particularly upon DNA damage. To verify that mitochondrial health is also affected in NER deficient animals, we performed the same test in *xpa-1(ok698)*. We observed a significant decrease in MMP following DNA damage induction (-49 %) suggesting that impaired mitochondrial function is triggered in response to unrepaired nuclear DNA damage **(Figure 5C)**.

**Figure 5.**
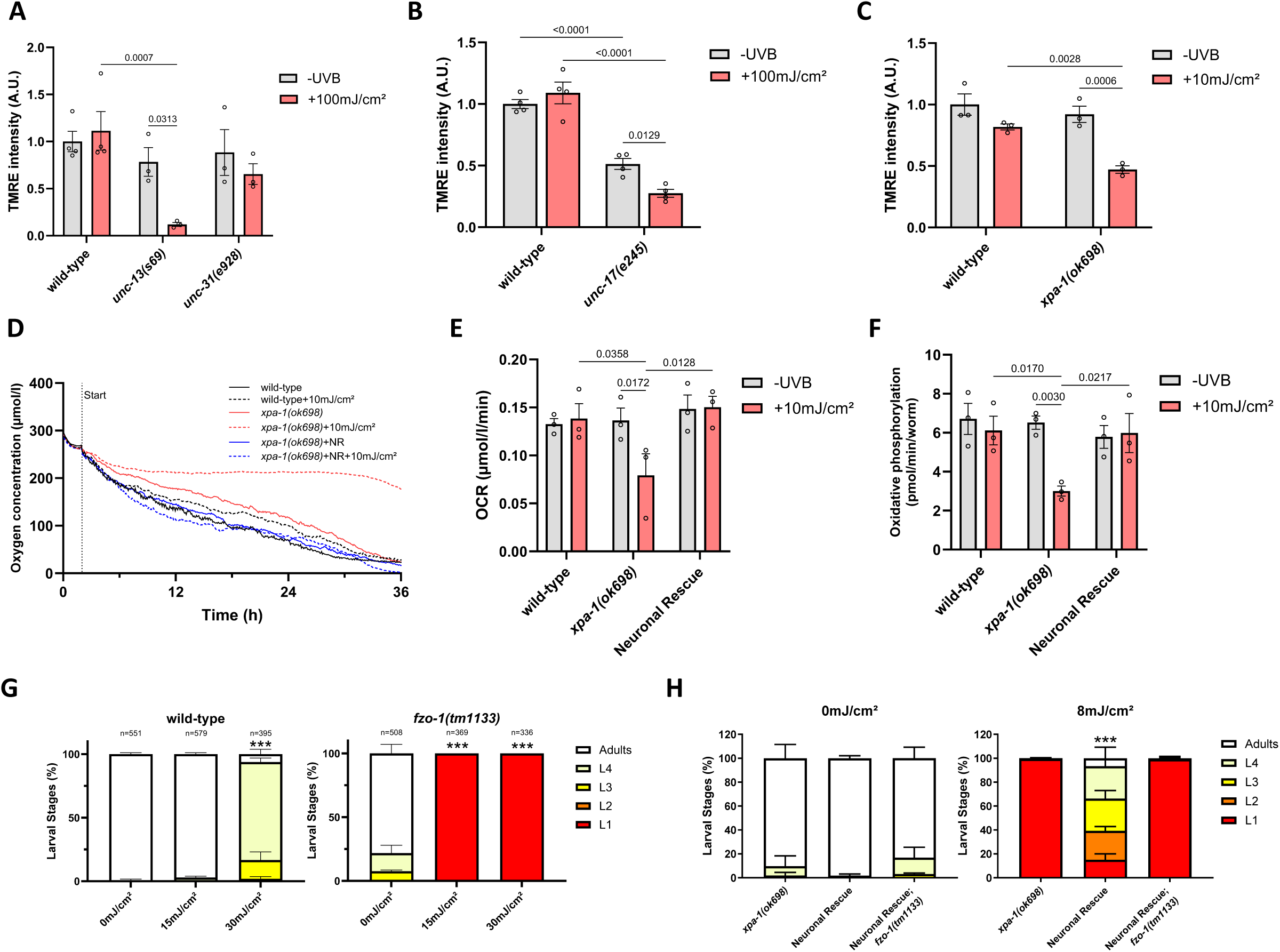
Mitochondrial activity mediates the systemic effect of neuronal *xpa-1* rescue. **(A)** Quantification of mitochondrial membrane potential using TMRE staining on day 1 adults from wild-type, neuropeptide deficient (*unc-13(s69)*) and neuroreceptor deficient (*unc-31(e928)*) strains, 24 hours after UVB irradiation. Data represent Mean+SD of N=3 biological replicates. Statistical analysis: Two-way ANOVA followed by Post-hoc Tukey. **(B)** Quantification of mitochondrial membrane potential using TMRE staining on day 1 adults from wild-type and Ach-signaling deficient strain *unc-17(e245)*, 24 hours after UVB irradiation. Data represent Mean+SD of N=3 biological replicates. Statistical analysis: Two-way ANOVA followed by Post-hoc Tukey. **(C)** Quantification of mitochondrial membrane potential using TMRE staining on day 1 adults from wild-type and *xpa-1(ok698)* strains, 24 hours after UVB irradiation. Data represent Mean+SD of N=3 biological replicates. Statistical analysis: Two-way ANOVA followed by Post-hoc Tukey. **(D)** Presense Oxodish measurment of oxygen concentration in wells seeded with day 1 adults from wild-type, *xpa-1(ok698)* and Neuronal rescue strains, 24 hours after UVB irradiation. Representative measurement from one biological replicate. Oxygen concentration was monitored during 36 hours starting 24 hours after UVB irradiation. **(E)** Quantification of Oxygen Consumption Rate from curves displayed in (D). Data represent Mean+SD of N=3 biological replicates. Statistical analysis: Two-way ANOVA followed by Post-hoc Tukey. **(F)** Oroboros measurement of phosphorylating respiration of day 1 adult stage wild-type, *xpa-1(ok698)* and Neuronal rescue strains, 24 hours after UVB irradiation. Data represent Mean+SD of N=3 biological replicates. Statistical analysis: Two-way ANOVA followed by Post-hoc Tukey. **(G)** Quantification of development of wild-type and mitochondrial fusion mutant *fzo-1(tm1133)* strain, 72 hours after UVB irradiation at L1 stage. Data represent Mean+SD of N=3 biological replicates. Statistical analysis: ANOVA. *p*-value:*<0.05, **<0.01, ***<0.001. **(H)** Quantification of development of NER deficient *xpa-1(ok698)*, Neuronal rescue and Neuronal rescue*;fzo-1(tm1133)* strains, 72 hours after UVB irradiation at L1 stage. Data represent Mean+SD of N=3 biological replicates. Statistical analysis: ANOVA. *p*-value:*<0.05, **<0.01, ***<0.001.

Next, we analyzed mitochondrial activity in wild-type, *xpa-1(ok698)* mutants and Neuronal rescue after UV exposure at Day 1 adult stage. To more directly evaluate mitochondrial activity, we made use of two devices measuring oxygen consumption. The first one, the Presens Oxysensor, allowed us to monitor the global oxygen consumption for a period spanning 36 hours following DNA damage induction. The second, the Oroboros system, allowed us to measure oxidative phosphorylation, 24 hours after DNA damage induction. Our Oxysensor measures showed that after UV exposure, *xpa-1(ok698)* mutants have a strongly decreased respiration rate compared to control worms (-71%). On the contrary, Neuronal rescue worms showed no significant difference in oxygen consumption **(Figure 5D, E)**. Accordingly, the oxidative phosphorylation in *xpa-1(ok698)* mutants after UV exposure was reduced (-54%) compared to wild-type, while Neuronal rescue animals showed no decrease **(Figure 5F)**. These results indicate that after DNA damage, NER activity in neurons is sufficient to maintain the whole organism’s mitochondrial activity.

Animals that are defective in mitochondrial fusion (*fzo-1(tm1133)*), a mechanism necessary for correct mitochondrial activity were recently observed to be hypersensitive to UV^42^. Accordingly, we observed a striking developmental delay after UV exposure in *fzo-1(tm1133)* mutants **(Figure 5G)**. We then ablated mitochondrial activity in the Neuronal rescue strain by generating the new strain: *fzo-1(tm1133);xpa-1(ok698)*;[*p_rgef-1_::gfp::xpa-1;p_unc-122_::gfp*] (i.e. *fzo-1(tm1133)*+Neuronal rescue). In this case, the efficiency of the Neuronal rescue to revert DNA damage sensitivity was completely ablated **(Figure 5H)**. These results establish the role of mitochondrial activity in the rescue of DNA damage sensitivity due to neuron-specific DNA repair.

Taken together, these data indicate that neuronal DNA repair is sufficient to maintain the organism’s mitochondrial activity through ACh signaling **(Figure 6)**.

**Figure 6.**
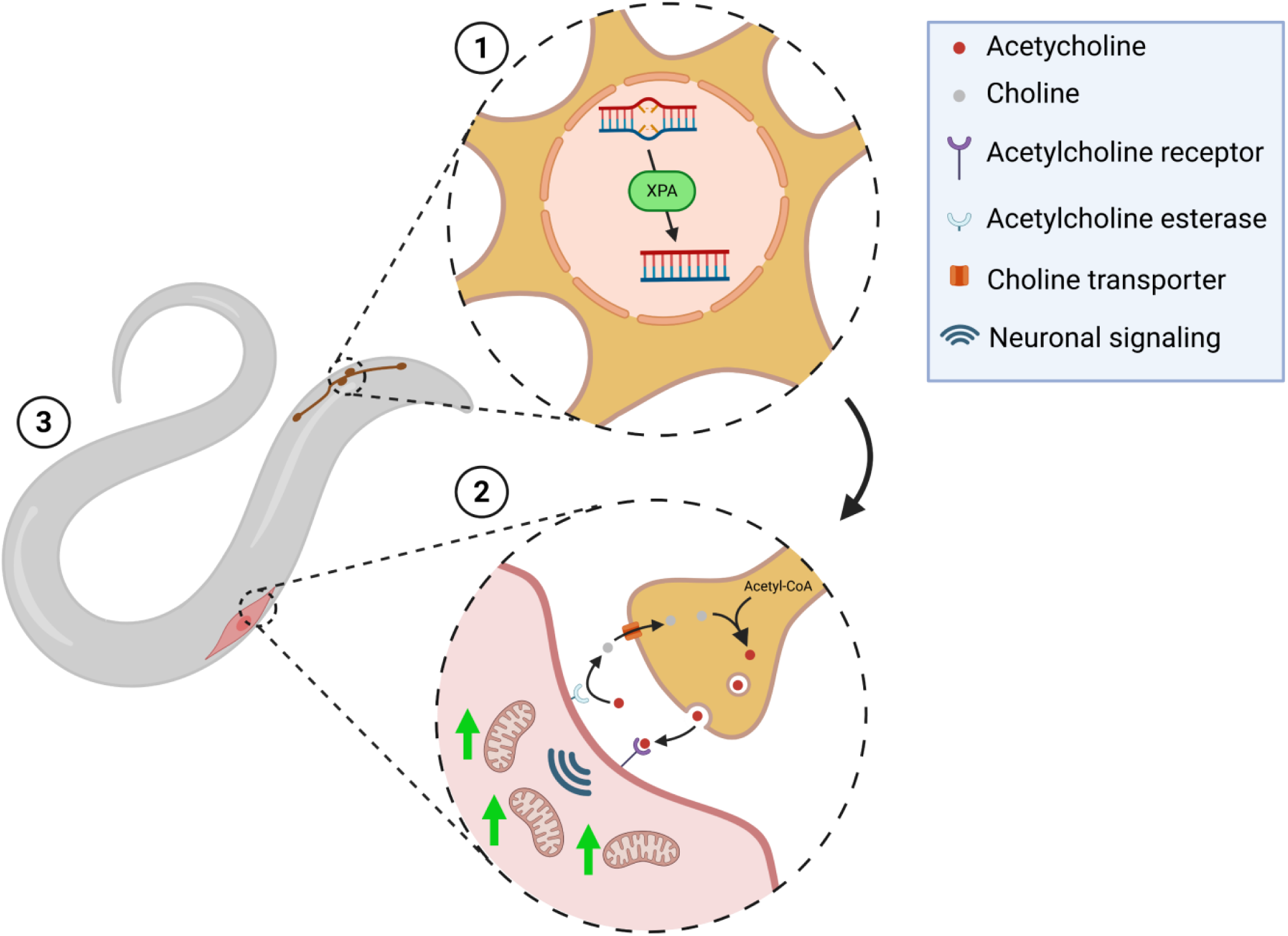
Proposed model. In wild-type *C. elegans*, bulky adducts are normally repaired by the Nucleotide Excision Repair pathway. In *xpa-1(ok698)* mutant strain, NER is fully ablated and bulky adducts remain in the genome, leading to genetic stress, decreased fitness and premature death. In our *xpa-1* re-expression strain (Neuronal rescue), we propose that ① bulky lesions are removed in neurons by the NER pathway allowing their survival. Thus ② maintaining neuronal signaling and more specifically acetylcholine signaling. Acetylcholine signaling maintenance allows the maintenance of high mitochondrial activity after UV irradiation and promotes muscle activity. We propose that ③ recovery of the neuromuscular axis promotes movement, development and ultimately rescues DNA damage sensitivity.

## Discussion

Thus far, DNA repair deficiency in specific organs has been shown to exert systemic pathological consequences. For example, hematopoietic NER deficiency in mice was shown to lead to cellular senescence in other, repair proficient cell types in mice^43^. Moreover, DNA damage-triggered dysfunction in specific organs can compromise the function of other organ systems, for example, chronic liver disease, often triggered by oxidative stress and inflammation^44^ was associated with multiple other pathologies^45^; and kidney insufficiency elevates a broad range of age-related disease risks as hypertension and diabetes^46,47^. Here, we report the opposite scenario: we probed the consequences of cell-type specific DNA repair recovery and observed that XPA-1 re-expression in either intestinal, muscle or neuronal cells can contribute to some degree to organismal resistance to DNA damage.

Strikingly, the repair of DNA lesions in neurons has the broadest effect on the organism’s ability to tolerate DNA damage. This could be explained by the maintenance of a tissue that is normally very sensitive. Indeed, mature neurons are described as sensitive to DNA damage due to general lower DNA repair efficiency^48,49^ and lower threshold for apoptosis^50^. Moreover, re-expression of the NER component XPF in a tissue-specific manner confirms that muscular cells are also less sensitive to DNA damage^51^. Here, we observed a clear and previously unknown effect of NER activity in neurons on the whole organism.

To achieve this resilience in peripheral tissue types, we determined that ACh signaling is essential. ACh signaling is an important component of neuronal signaling^52^ that among a range of functions sets the capacity to tolerate protein folding stress^53^ and impacts mitochondrial stress-response^54^. Here we show that loss of ACh signaling leads to a strong DNA damage sensitivity. Little is yet known about the link between ACh signaling and DNA repair. Loss of brain nicotinic acetylcholine receptor is one of the hallmarks of neurodegenerative disorders as Alzheimer’s disease ^55^ and decreased oxidative DNA damage has been proposed as the mechanism of action of Memantine, a drug used against Alzheimer’s disease^56^. It is tempting to speculate that the effect of Memantine might allow the maintenance of ACh signaling and potentially mitochondrial activity in a non-cell autonomous manner thus benefiting cells beyond cholinergic neurons.

Growing evidence shows that the DNA damage response and mitochondrial activity are interconnected. It was proposed that mitochondrial NAD^+^ reserves are deprived and Sirtuins levels decreased amid persistent DNA lesions^57^, resulting in decreased PGC1α and FOXOs levels leading to altered mitochondrial morphology^58,59^. Vice versa, dysfunctional mitochondrial dynamics also leads to increased DNA damage sensitivity^42^. Here, we demonstrated that the recovery of NER in neurons leads to maintenance of mitochondrial activity after UV-induced DNA damage in peripheral tissues. This shows for the first time that alterations in the genome maintenance of one cell not only impacts the mitochondria of the affected cell but also mitochondria in other cells. As mitochondria can operate as a hub for many different signaling pathways^60,61^, it will be interesting to further explore the organismal consequences of restored mitochondrial activity amid DNA damage accumulation.

The mechanism by which ACh maintains mitochondrial activity after DNA damage remains unexplored. Mitochondria have the capacity to sense ACh, notably as they express several nicotinic acetylcholine receptor^62^ and activation of such receptors prevents mtDNA release^63^. Moreover, impairment of ACh signaling leads to increase in mtUPS^54^. However, our results show that treating the animals with ACh is not sufficient to mimic DNA repair effects in neurons potentially due to inadequacy of extrinsic supplementation to affect the ACh availability at a synapsis. Indeed, the loss of acetylcholine esterase did not lead to DNA damage sensitivity indicating that, contrary to ACh *per se*, it is the maintenance of the neuronal ACh influx that leads to maintenance of mitochondrial activity in other tissues.

We thus propose that neuronal DNA repair is decisive for *C. elegans* overall susceptibility to DNA damage. The organismal response to DNA damage critically depends on mitochondrial activity, which can be modulated via neuronal ACh signaling. Our results for the first time establish that not only cell type specific DNA repair deficiency but also cell type specific DNA repair determines the organism’s ability to grow and survive amid DNA damage infliction.

## Methods

### *C. elegans* strains and maintenance

*C. elegans* worms were grown according to standard practices^64^. In short, worms were kept at 20°C on NGM plates seeded with *E.coli* strain OP50 and kept in sealed boxes in light-protected incubators. All strains used in this study are summarized in Supplementary Table S1.

### Generation of transgenic lines

Transgenic *C. elegans* lines were generated following previously described protocol^65^. To generate the tissue specific XPA-1::GFP expression strains, each specific promoter was amplified by PCR (2kb upstream respective ATG; Pxpa-1: FW: ACTAAGCTTGCACCAATTGGTGCATGTTC, RV: AGTGTCGACCGGCATTTATCTGAAACAATCG; Pmyo-3: FW: ATAGCGGCCGCTTCTAGATGGATCTAGTGG, RV: TATGGCGCGCCGATTGCTTTTTCACAATCAG; Pges-1: FW: ATAGCGGCCGCCTGAATTCAAAGATAAGATATG, RV: TATGGCGCGCCCTCCGAACTATGATGAC; Prgef-1: FW: ATAGCGGCCGCCGTCGTCGTCGTCGATG, RV: TATGGCGCGCCGCTCAATTGAATGACGTCACAGG) and inserted in pPD95 worm expression plasmid. *xpa-1* gene was then amplified by PCR on wild-type DNA and inserted in resulting plasmids. Young adults of the RB864 strain ((*xpa-1(ok896*)) were injected using a microinjection system and pCFJ90 (*pmyo-2::tdTomato]*) or pCFJ68 (*punc-122::gfp*) were used as co-injection markers. Extrachromosomal arrays were integrated using gamma irradiation and backcrossed 4 times.

### Hypochlorite synchronization

Gravid adult worms were washed with 10mL of 1x M9 buffer and collected. The worms were centrifuged 1min at 350g and the supernatant discarded. The pellet was resuspended in 3mL of alkaline hypochlorite solution (10% Sodium hypochlorite, 5 M NaOH) and incubated 3min with gentle shaking. The reaction was stopped by addiction of 10ml M9 buffer. The remining eggs were centrifuged 2min at 800g and the supernatant removed. The eggs were washed 3 times in 5ml M9 buffer then incubated overnight in 5ml of M9 Buffer on a rotator (RM10W-80V CAT). Next day, prior to seeding the L1 larvae were filtered on a 11 µM hydrophilic filter (Merck Millipore) concentrated and counted.

### UV irradiation

For L1 larvae, after counting, the appropriate number of worms was seeded on NGM agar plates freshly seeded with OP50 bacteria to avoid shielding of the UVB. The worms were let on bacteria for 4 hours to recover from overnight starvation. The worms were then placed under a UVB lamp (Waldmann Medizinthechnik UV 236 B) for the appropriate time with lead open to avoid shielding of the UVB.

For Day 1 adult worms, synchronized plates were washed with 10mL of 1x M9 buffer and collected. The worms were centrifugated 1min at 350g. The supernatant was discarded leaving a few hundred microliter to transfer the worms on a non-seeded NGM plate to avoid shielding of the UVB. The plates were let to dry under a fume hood for several minutes and placed under a UVB lamp (Waldmann Medizinthechnik UV 236 B)) for the appropriate time with lead open to avoid shielding of the UVB. The worms were then washed with 10ml M9 buffer, centrifugated 1min at 350g, the supernatant was discarded leaving a few hundred microliter and the worms transferred on a OP50 seeded NGM plate. The power of the lamp was measured using a AnalytikJena UVX Radiometer and the exposure time was calculated as follows:

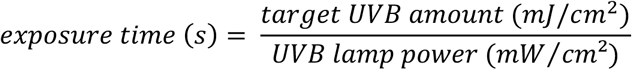

The amount of UVB was adapted to the worms developmental stage and the DNA damage sensitivity of each strain with a range of 0mJ/cm² to 10mJ/cm² on L1 worms (0 to 4 with *xpa-1(ok698)*) and 0mJ/cm² to 100mJ/cm² on Day 1 adult worms (0 to 25 with *xpa-1(ok698)*).

### Developmental assay

Hypochlorite synchronization was performed and around 200 L1 worms were UVB irradiated. The worms were then incubated 72 hours at 20°C. Once wild-type worms reached adulthood, plates were imaged at 30x magnification using a Zeiss Axio Zoom. V16. Worms were categorized as L1, L2, L3, L4 and adults using the ImageJ software and plotted as percentage of the whole population.

### Lifespan assay

Synchronized populations of *C. elegans* worms (60 to 90 worms per condition) were irradiated at day 1 adulthood and transferred into OP50 seeded NGM plates. The worms were then maintained at 20°C and monitored every day for dead worms. Worms found dried on the walls of the plates, dead due to internal hatching or during transfer were censored. To avoid progeny accumulation the worms were transferred to new plates everyday as long as laying eggs. Results were then plotted using the “Survival” option of the GraphPad Prism 10 software.

### Phalloidin staining

Phalloidin staining experiments were performed as previously described^21^, with some modifications. Prior to the staining procedure, a phalloidin-iFluor stock solution (1000x Phalloidin-iFluor 647 reagent (Abcam) in DMSO) was prepared and kept at-20°C. Synchronized populations of *C. elegans* worms were UVB irradiated at day 1 adulthood. 72h after irradiation, worms were collected and washed with M9 buffer and fixed with 4% paraformaldehyde solution for 15-20 mins. Worms were then permeabilized with 2% Tween-20 in 1xPBS for 30 mins at room temperature (RT) and incubated in ß-mercaptoethanol solution (120mM Tris-HCl; 5% ß-mercaptoethanol; 1% Triton X-100) for 10 mins at RT. Afterwards, worms were washed 3x with 0.2% 1xPBST and 3x with 1% BSA in 0.2% 1xPBST. Worms were stained with a freshly prepared 2x phalloidin working solution (2% phalloidin-iFluor stock solution in 1xPBS) for 1h at RT, covered from light, on a rotator (RM10W-80V CAT). Worms were then washed 3x with 0.2% 1xPBST, transferred to microscope slides and cover slips were mounted using FluorSave Reagent (Merck). Worms were imaged with a Zeiss AxioImager M1 microscope at 40× magnification.

### Motility assay

Synchronized populations of *C. elegans* worms were irradiated with UVB doses at day 1 adulthood. 72h after irradiation, 30-40 worms were transferred to non-seeded NGM agar plates and filmed, 30min later, for 30s using a Zeiss AXIO Zoom.V16 stereomicroscope. Video analysis, and quantification of average and maximum speed values were done using the “wrMTrck” ImageJ plugin as previously described^66^.

### Pharyngeal pumping assay

Synchronized populations of *C. elegans* worms were irradiated with UVB at day 1 adulthood. 24h after, the number of pumps in 30s was manually counted using a clicker and a Zeiss AXIO Zoom.V16 stereomicroscope.

### DiI filling

Prior to the staining procedure, a DiI stock solution (2mg/mL 1,1’-dioctadecyl-3,3,3’,3’-tetramethylindocarbocyanine perchlorate (DiI) in dimethyl formamide) was prepared and kept at-20°C. Synchronized populations of *C. elegans* worms were irradiated UVB at day 1 adulthood. 48h after irradiation, worms were collected from the plates and washed 3 times with 1x M9 buffer. Afterwards, worms were incubated in 5 mL of freshly prepared DiI staining solution (1:200 dilution of DiI stock solution in 1x M9) for 45 min, in the dark, at RT, on a rotator (RM10W-80V CAT). Worms were then centrifuged (1min, 350g) and washed 3 times with M9 buffer. Excessive staining was remove by letting the worms 1 hour on heat inactivated (2hours at 65°C) OP50 seeded plates for 1hour. Afterwards, 15- 20 worms were transferred to 2% agarose pads in microscope slides via picking, anesthetized with 5 mM Levamisole (Levamisol(−)-Tetramisol-hydrochlorid, L9756-5G Sigma-Aldrich) and imaged with a Zeiss AxioImager M1 microscope with 10× magnification. Exposure time was selected for each experiment based on the fluorescence signal of the non-irradiated wild-type population. Worms with visible stained phasmid cell bodies were manually quantified.

### Intestinal barrier function assay

Intestinal barrier function assays (Smurf assays) were performed as previously described^67^, with some modifications. Synchronized populations of *C. elegans* worms were irradiated with UVB at day 1 adulthood. 72h after irradiation, worms were washed 1x with M9 buffer and incubated in 500 µL Blue Food Solution (50 µL 5% Brilliant Blue FCF (Sigma-Aldrich) in H_2_O; 50 µL 3x concentrated *E. coli* OP50 overnight culture in LB medium and 400 µL M9 buffer) for 6h at RT on a rotator (RM10W-80V CAT). Worms were then washed once with M9 buffer, collected by gravity settling on ice and transferred into empty NGM agar and let to dry under a fume hood. Finally, worms were transferred to 2% agarose pads in microscope slides via picking, anesthetized with 5 mM Levamisole (Levamisol(−)-Tetramisol-hydrochlorid, L9756-5G Sigma-Aldrich) and imaged with a Zeiss AxioImager M1 microscope with 5× magnification. Exposure time was selected for each experiment based on the fluorescence signal of the non-irradiated wild-type population. Worms with visible dye spread all over the body cavity (Smurf phenotype) were manually quantified.

### RT-qPCR

Synchronized population of 2000 Day 1 adults animals were UV irradiated. 24 hours after, the animals were washed three time in 1x M9 buffer before snap freeze in 1ml TRIzol (Invitrogen, 15596018). Total RNA was extracted using the RNeasy Mini Kit (QIAGEN, 74106) following manufacturers guide but replacing the chlorophorm by 1-bromo-3-chloropropane (Sigma, B9673). RNA quantity was quantified using NanoDrop 8000 (Thermo Fisher Scientific, ND-8000-GL) and quality tested by assessing ribosomal RNA on a electrophoresis gel. 1µg of RNA was reverse transcribed using the Superscript III (Invitrogen, 18080044) and qPCR was performed in a BIO-RAD CFX96 real-time PCR machine (BIO-RAD, 1855196) using PowerUp SYBR Green Master Mix (Applied Biosystem A25742). Quantity of *cha-1* expression was normalized to three different housekeeping genes: *eif3.c, tbg-1* and *act-1*. Primer sequences: *cha-1* FW: CGTGGTGTGTCTTGACATGGA, RV: CGATTGAGCCCGAACTTTGA. *eif-3.c* FW: ACACTTGACGAGCCCACCGAC, RV: TGCCGCTCGTTCCTTCCTGG. *tbg-1* FW: CAATGTGCCCATCAATTCGG, RV: AACAAGAAGCGAGTGACGTC. act-1 FW: CCACCATGTACCCAGGAATT, RV: AGAGGGAAGCGAGGATAGAT. Relative mRNA levels were quantified using the 2^-ΔΔCt^ method^68^.

### TMRE staining

TMRE staining experiments were performed as previously described^58^. Synchronized populations of *C. elegans* worms were irradiated with UVB at day 1 adulthood. 24h after irradiation, worms were transferred on NGM agar plates, previously seeded with 30 µM TMRE (Tetramethylrhodamin-Ethylester-Perchlorat 87917-25MG Sigma Aldrich) dissolved in heat-inactivated OP50 (2hours at 65°C). The worms were incubated for 2hours before to be transferred to heat inactivated OP50 seeded NGM plates to remove extra dye from the intestine. Then, 20-30 worms were transferred to 2% agarose pads in microscope slides via picking, anesthetized with 5 mM Levamisole (Levamisol(−)-Tetramisol-hydrochlorid, L9756-5G Sigma-Aldrich) and imaged with a Zeiss AxioImager M1 microscope with 5× magnification. For the analysis, mean fluorescence of the whole worm body was quantified using ImageJ.

### Oxygen consumption measurement

SDR SensorDish® Reader: Synchronized populations of *C. elegans* worms were irradiated with UVB at day 1 adulthood. 24h after irradiation, worms were washed 5 times with M9 buffer to remove bacteria and resuspended in complete liquid NGM media mixed with inactivated OP50 (2hours at 65°C, OD=0.8). 100 worms were platted in 6ml NGM per well in a DeepWell OxoDish. The plate was sealed with a BioRad Microseal® Adhesive Sealer (MSB-001) and incubated 20°C. The oxygen concentration in each well as measure every minute for 36 hours using the SDR SensorDish® Reader and the SDR 4.0 software from the Presens company. Quantification was made by calculating the slope of for each curve at the linear state.

Oroboros: Synchronized populations of *C. elegans* worms were irradiated with UVB at day 1 adulthood. 24h after irradiation, worms were washed 5 times with M9 buffer to remove bacteria and resuspended in complete liquid NGM media. After calibration the worms were transferred in the Oroboros O2K-oxygraph. The oxygen flux was measured before (Basal respiration) and after addition of 10µl of 5M Sodium-Azide (S2002 – Sigma Aldrich) (Non-mitochondrial respiration) and the phosphorylating respiration rate was calculated by subtracting the Non-mitochondrial respiration from the Basal respiration.

### Statistics

Statistical analysis was performed using the GraphPad Prism 10 software package (GraphPad Software Inc., San Diego, USA). The statistical tests applied for each experiment are reported in the figure legends. Only *p*-value of significant differences (<0.05) are displayed on graphs. For developmental assays, “n=” represent the total number of animals scored among all biological replicates. For developmental assays, a modified ANOVA was performed using the following definitions for the total sum of squares (*SSt*) and the within group sum of squares (*SSw*), with *g* being groups, 𝑛 the number of replicates and 𝑥 the vector of ratios of the stages:

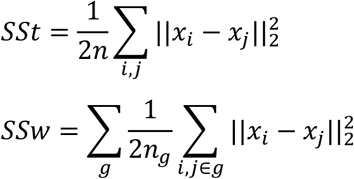

Unless stated, the test was performed against the control condition. For readability, the significance was displayed as follow: *=*p*<0.05; **=*p*<0.01; ***=*p*<0.001. Exact *p*-values can be found in Supplementary Table 2.

## Data availability

Original data are provided in the source data file.

## Acknowledgments

Nematode strains were provided by the National Bioresource Project (supported by The Ministry of Education, Culture, Sports, Science and Technology, Japan), the Caenorhabditis Genetics Center (funded by the NIH National Center for Research Resources, US), and the *C. elegans* Gene Knockout Project at the Oklahoma Medical Research Foundation (part of the International *C. elegans* Gene Knockout Consortium). We thank Walter Sandt for providing help for statistical analysis.

F.L.d.S. received support from the Cologne Graduate School of Ageing Research. B.S. acknowledges funding from the European Research Council (ERC-2023-SyG, 101118919), the Hevolution Foundation (HF-GRO-23-1199212-35), the Deutsche Forschungsgemeinschaft (Reinhart Koselleck-Project 524088035, FOR 5504 project 496650118, FOR 5762 project 531902955, SFB 1678, SFB 1607, CECAD EXC 2030 – 390661388, DFG-ISF project 561031107, ANR-DFG project 545378328, and the DFG project grants 558166204, 540136447, 496914708, 437825591, 437407415, 418036758), the José Carreras Leukemia Foundation, DJCLS 04 R/2023, the Deutsche Krebshilfe (70114555), and the John Templeton Foundation Grant (61734). The final model was generated using Biorender.

## Author contributions

P.S., B.R., M.R., C.R. performed and analyzed experiments; P.S., B.R., M.R. designed the experiments; P.S., M.R., B.S. conceived the study; P.S., B.R., B.S. wrote the manuscript; B.S. acquired the funding.

---

**Supplementary Figure 1.**
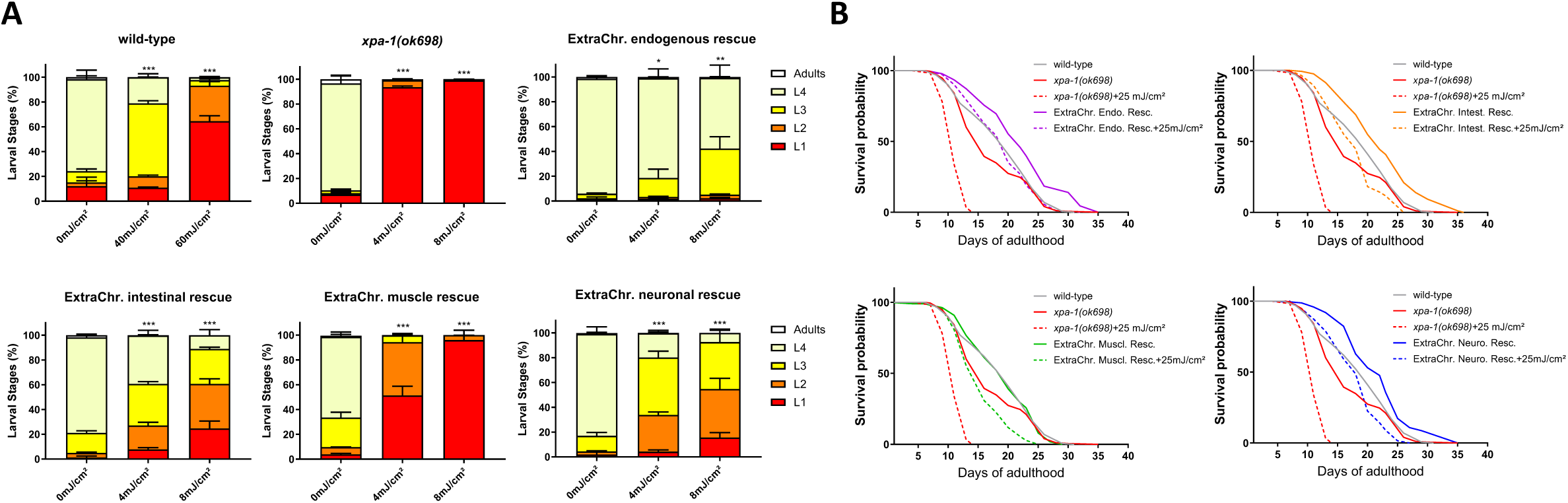
Extrachromosomal XPA-1::GFP expression rescues developmental arrest upon. (**A**) Quantification of development of wild-type, *xpa-1(ok698)* and extrachromosomal re-expression of XPA-1 strains, 72 hours after UVB irradiation at L1 stage. Data represent Mean+SD of N=3 biological replicates. Statistical analysis: ANOVA. *p*-value:*<0.05, **<0.01, ***<0.001. (**B**) Lifespan assays of extrachromosomal re-expression of XPA-1 strains after UVB irradiation at day 1 adult stage. Representative survival curve from one individual experiment. n>100 per condition.

**Supplementary Figure 2.**
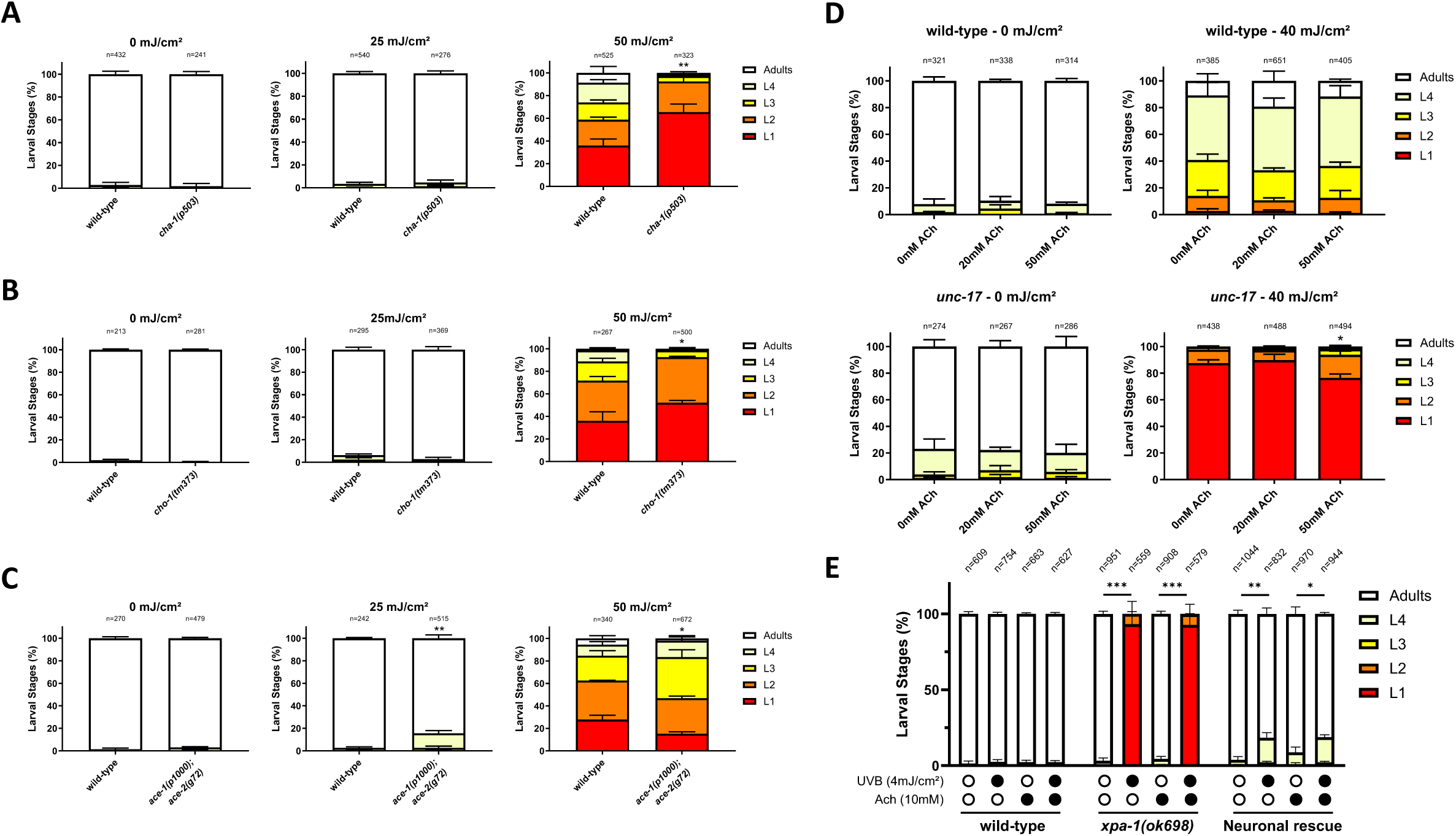
Ach signaling impairment leads to DNA damage sensitivity but is not rescued by Ach supplementation. **(A)** Quantification of development of wild-type and acetylcholine signaling mutant *cha-1(p503)* strains, 72 hours after UVB irradiation at L1 stage. Data represent Mean+SD of N=3 biological replicates. Statistical analysis: ANOVA. *p*-value:*<0.05, **<0.01, ***<0.001. **(B)** Quantification of development of wild-type and acetylcholine signaling mutant *cho-1(tm373)* strains, 72 hours after UVB irradiation at L1 stage. Data represent Mean+SD of N=3 biological replicates. Statistical analysis: ANOVA. *p*-value:*<0.05, **<0.01, ***<0.001. **(C)** Quantification of development of wild-type and acetylcholine signaling mutant *ace-1(p1000);ace-2(g72)* strains, 72 hours after UVB irradiation at L1 stage. Data represent Mean+SD of N=3 biological replicates. Statistical analysis: ANOVA. *p*-value:*<0.05, **<0.01, ***<0.001. **(D)** Quantification of development of wild-type and acetylcholine signaling mutant *unc-17(e245)* strains treated with 0, 20 or 50mM acetylcholine, 72 hours after UVB irradiation at L1 stage. Data represent Mean+SD of N=3 biological replicates. Statistical analysis: ANOVA. *p*-value:*<0.05, **<0.01, ***<0.001. **(E)** Quantification of development of wild-type, NER deficient *xpa-1(ok698)* and Neuronal rescue strains treated with 10mM acetylcholine, 72 hours after UVB irradiation at L1 stage. Data represent Mean+SD of N=3 biological replicates. Statistical analysis: ANOVA. *p*-value:*<0.05, **<0.01, ***<0.001.

